# Integrating defense and leaf economic spectrum traits in a tropical savanna plant

**DOI:** 10.1101/2022.12.11.519980

**Authors:** Neha Mohanbabu, Michiel P. Veldhuis, Dana Jung, Mark E Ritchie

## Abstract

1. Allocation to plant defense traits likely depends on resource supply, herbivory, and other plant functional traits such as the leaf economic spectrum (LES) traits. Yet, attempts to integrate defense and resource acquisitive traits remains elusive.
2. We assessed intraspecific correlations between different defense and LES traits in a widely distributed tropical savanna herb, *Solanum incanum*, a unique model species for studying allocations to physical, chemical, and structural defenses to mammalian herbivory.
3. In a multivariate trait space, the structural defenses - lignin and cellulose - were positively related to the resource conservative traits - low SLA and low leaf N. Phenolic content, a chemical defense, was positively associated with resource acquisitive traits - high SLA and high leaf N - while also being associated with an independent third component axis. Both principal components 1 and 3 were not associated with resource supply and herbivory intensity. In contrast, spine density - a physical defense - was orthogonal to the LES axis and positively associated with soil P and herbivory intensity.
4. Synthesis: These results suggest a hypothesized “pyramid” of trade-offs in allocation to defense along the LES and herbivory intensity axes. Therefore, future attempts to integrate defense traits with the broader plant functional trait framework needs a multifaceted approach that accounts for unique influences of resource acquisitive traits and herbivory intensity.

## Introduction

Plant traits are strongly affected by both resource availability and biotic interactions. The key traits of the leaf economic spectrum (LES) (Reich et al. 1999, Wright et al. 2004, Westoby and Wright 2006, Donovan et al. 2011), which encompass traits from conservative to acquisitive resource-use strategies, are associated with different resource supplies in the environment. Low resource supplies are associated with conservative strategies such as slow growth rate, low photosynthetic rate and long leaf lifespan and high resource supplies are associated with acquisitive strategies such as fast growth rate, high photosynthetic rate and short leaf lifespan. Likewise, plant traits that defend against herbivory (defense traits such as spines and secondary metabolites) may be similarly affected by resource supply and may correlate with the LES axis (Coley et al. 1985, Herms and Mattson 1992). Consequently, certain defense traits are likely to co-occur with specific LES traits resulting in trait syndromes (Defossez et al. 2018, Agrawal 2020, Armani et al. 2020). Studying such trait syndromes are key to understanding whole plant strategies under complex natural environments (Pellissier et al. 2018, Blumenthal et al. 2020) for both within-species and across species comparisons. However, attempts to study intraspecific covariation in defense and LES traits are rare, especially for plants growing under natural conditions that simultaneously vary in multiple resources.

Plant defense and LES traits may be correlated in several different ways depending on associations between resource supply and risk from herbivory. For instance, at high resource supply, resource acquisitive traits such as high specific leaf area (SLA), high leaf nitrogen (N) and short leaf lifespan (LLS) create high quality tissue, which may also make the plants more vulnerable to herbivory; thus, these plants may invest in secondary defensive metabolites (metabolic defense strategy) (Chauvin et al. 2018, Agrawal 2020, Morrow et al. 2022) (Fig 1a,b). In contrast, at low resource supply resource conservative traits such as low SLA, low leaf N (i.e., low plant quality) and long LLS may also deter herbivores or support low herbivore abundance, likely causing structural defense traits such as cellulose and lignin to be negatively associated with resource acquisitiveness (Mason and Donovan 2015, Abdala-Roberts et al. 2018, Agrawal 2020) (Fig 1a,d) (avoidance strategy). Taken together, these expectations are similar to the predictions of the Resource Availability Hypothesis *(sensu* Coley et al. 1985) which suggests that species in high resource environments are likely to invest in inducible chemical defenses whereas species in low resource environments should invest in constitutive or structural defenses. More recently, Armani et al. (2020) extend these results to physical defense traits by showing a negative association between a structural defense, thorn mass fraction, and SLA. Alternatively, if herbivory risk is independent of the resources driving LES traits and/or predation risk or thermal conditions in the ecosystem (Ritchie 2000, Anderson et al. 2010, Veldhuis et al. 2020), (Fig 1a,c) then defense traits may be uncorrelated with resource acquisitiveness. Thus, there can be a complicated interaction between resource supply and herbivory that may influence both plant defense and LES traits.

**Figure 1:**
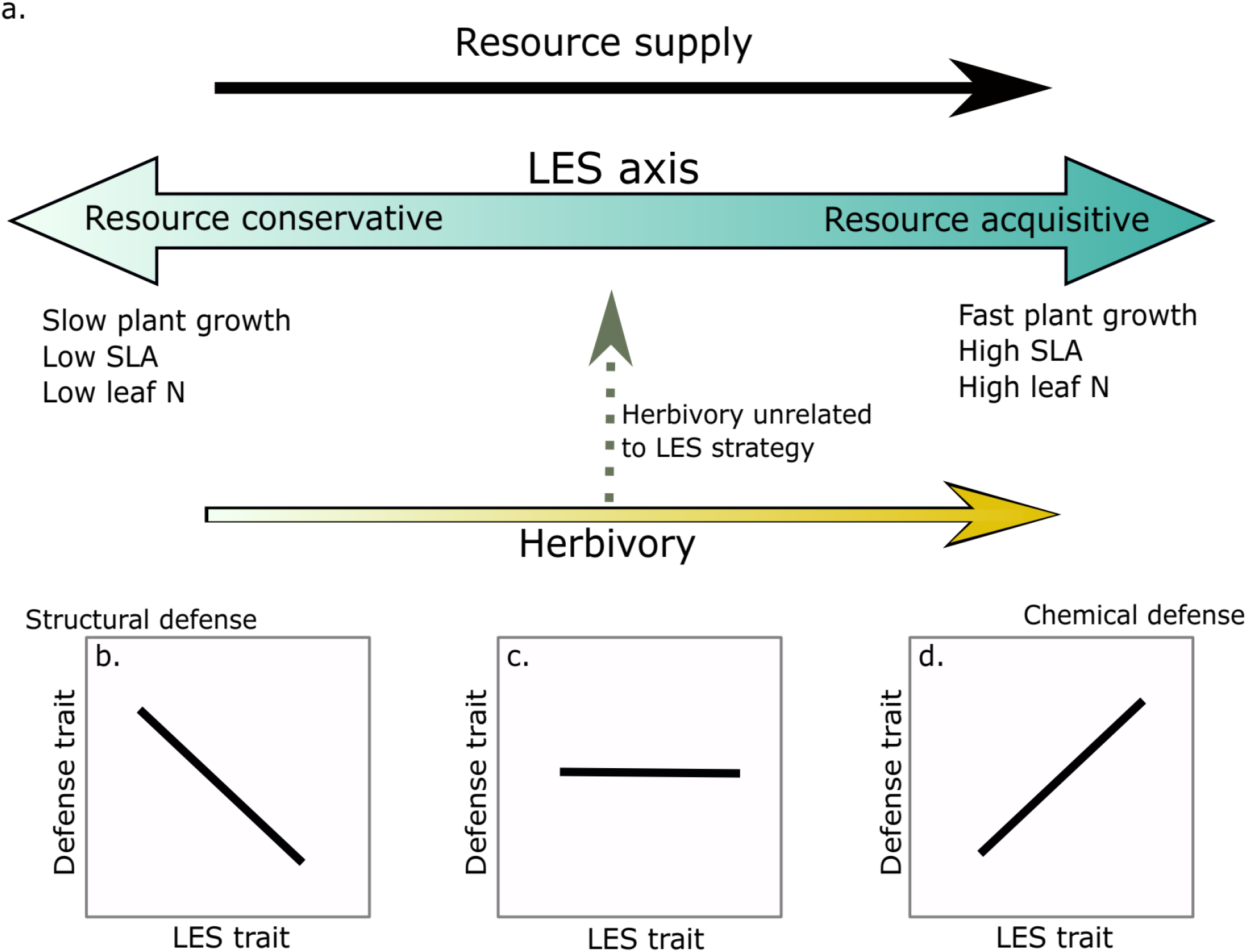
Potential associations between defense and LES traits for within species comparisons. a) The blue arrow gradient represents the LES axis. Light blue indicates a resource conservative strategy and darker blue a resource acquisitive strategy. Resource supply is assumed to be positively associated with resource acquisitiveness (black arrow). Risk from herbivory can be parallel to the LES axis (yellow arrow) or can be orthogonal to it (dashed grey arrow). b) Leaf structural defenses are negatively associated with LES at the conservative end of the spectrum; c) Lack of association between defense and LES traits when herbivory is unassociated with LES axis; and d) Chemical defense is positively associated with the LES traits at the acquisitive end of the spectrum.

Thus far, LES-defense trait associations have been exclusively tested along gradients of resources that increase both plant growth (i.e., promoting resource acquisitive strategy) and herbivory intensity (Defossez et al. 2018). But resource-herbivory relationships need not always be positive as evident from examples of decreasing herbivory in high resource environments or lack of association between resource and herbivory (La Pierre and Smith 2016, Mohanbabu and Ritchie 2022). Alternatively, herbivory maybe more strongly associated with resources different from those influencing plant growth strategy (e.g., sodium [Na]; Kaspari 2020, Prather et al. 2020). For instance, LES is generally associated with leaf N, and leaf N would be expected to increase with environmental N availability (Bracken et al. 2015), consequently defenses might decrease overall or shift from structural and physical to chemical with increasing N availability. However, herbivore abundance and intensity may respond to resources other than N, such as P or Na (Bishop et al. 2010, La Pierre and Smith 2016, Kaspari 2020, Prather et al. 2020, Mohanbabu and Ritchie 2022). If so, defenses may change more strongly in response to these other resources instead of N and consequently may not be strongly related to the LES. As both resource supply and herbivory can influence allocation to traits, studying trait associations along multiple resource gradients provides an opportunity to understand more complex trade-offs that are unlikely to occur in response to single resource gradients.

Furthermore, trade-offs in defense and LES traits across multiple species may be influenced by phylogenetic constraints that may limit certain combinations of traits (Eichenberg et al. 2015, Zhou et al. 2022). Exploring trait covariation within species provides an opportunity to study trade-offs in traits that are more likely to be an outcome of response to abiotic and biotic environments rather than a long evolutionary history. Both LES and defense traits show considerable intraspecific variation usually along gradients of climate, resources, and/or herbivory risk (Albert et al. 2010, Violle et al. 2012, Siefert et al. 2015, Hahn and Maron 2016, Fajardo and Siefert 2018, Moreira et al. 2018, Hahn et al. 2018, Lynn and Fridley 2019). Some recent work from two disparate systems-*Populus tremuloides* in temperate region (Morrow et al. 2022) and an invasive species, *Chromoleana odorata*, from the tropics (Li et al. 2022) indicates potential for trait covariation within species such that acquisitive plants are also better defended. But intraspecific covariation in defense and LES traits remains poorly understood in plants that experience mammalian herbivory.

Mammalian herbivores may impose very different cost-benefit constraints for defenses compared to insect herbivores consequently influencing traits that impact growth-defense trade-offs (Perkovich and Ward 2022). For example, mammalian herbivores may not be as sensitive to chemical defenses as insect herbivores due to their ability to neutralize chemical defenses by consuming a diverse group of plants (Mattson 1980). Under such a scenario, resource acquisitive plants may benefit from a tolerance strategy (i.e., no defense but high capacity for regrowth) rather than a chemical defense strategy. Alternatively, resource-conservative plants which have low plant quality due to elevated levels of lignin and cellulose may not deter large-bodied mammalian herbivores which can compensate for low quality food by increasing the amount of plants consumed (Olff et al. 2002), essentially reducing the efficacy of an avoidance strategy. Thus, plants experiencing considerable mammalian herbivory may optimize different combinations of LES and defense traits.

In this study, we explored intraspecific associations between LES and defense traits in *Solanum incanum* along gradients of rainfall, total soil N, and soil P in the Serengeti National Park, Tanzania. Our species of interest, *S. incanum*, serves as a unique model species for exploring the effect of mammalian herbivory on plant traits. It is widely distributed and produces different types of carbon-based defenses, including chemical defenses such as phenolics, leaf structural defenses (lignin and cellulose), and physical defenses (spines). Furthermore, members of the Solanaceae family have been well-studied in the context of chemical defenses against insect herbivores, similar to other model species for studying tradeoffs between plant defense and LES traits, such as *Asclepias* (Agrawal and Fishbein 2006, Züst et al. 2015, Agrawal 2020) and *Helianthus* (Mason and Donovan 2015, Mason et al. 2016). More recently, studies have used *Solanum sp*. to understand responses of physical defenses to mammalian herbivory and neighborhood interactions in the field (Coverdale et al. 2018, 2019). This diversity of defense types allows for specific comparisons in associations of different types of defense traits with resources and LES traits. We specifically asked: 1) Do defense and LES traits show bivariate correlation? 2) How are the different defense and LES traits correlated in a multivariate space? 3) Do resource availability and herbivory intensity (i.e., herbivory risk) explain associations between LES and defense traits?

## Study site and Methods

### Field site

Serengeti National Park sits within one of the last remaining grazing ecosystems in the world. Like other grasslands in East Africa, plants in the Serengeti have coevolved for millions of years with a diverse assemblage of resident and migratory herbivores such as wildebeest (*Connochaetes taurinus*), plains zebra (*Equus quagga*), Thomson’s and Grant’s gazelles (*Eudorcas thomsonii, Nanger soemmerringii*), impala (*Aepyceros melampus*), and elephant (*Loxodonta africana*) (McNaughton 1985). Therefore, the plants have likely evolved adaptations to survive under intense herbivory pressure with annual aboveground biomass losses often ranging from 60-90% (McNaughton 1985). This park also has natural gradients of rainfall (700-1125mm), soil N (0.2-3.7mgg^-1^) and soil P (0.007-0.53mgg^-1^) (Ruess and Seagle 1994, Anderson et al. 2007b).

### Study organism

*Solanum incanum* (hereafter *Solanum*), is a pan-African and pan-Asian herbaceous plant that is common and widely distributed in Serengeti. *Solanum* invests in both physical, structural, and chemical defenses making it a good “model” system to study variation in defense traits. Among the physical defenses, *Solanum* stems may be covered with spines that vary in density and can also be induced by herbivory (Coverdale et al. 2019). Additionally, *Solanum*, like other plants, may also allocate resources to leaf structural content such as lignin and cellulose which can have anti-herbivore defensive properties. Lastly, members of the Solanaceae have been reported to allocate resources to secondary defensive metabolites such as alkaloids and phenolics (Chowański et al. 2016, Kaunda and Zhang 2019).

### Study Design

The trans-Serengeti plots, first surveyed in the early 2000s, are a network of sites that were chosen from a selection of random points in 10 x 10 km grids spanning the geographical extent of the Serengeti (Anderson et al. 2007a). We revisited 61 of these plots in 2018 and found *Solanum* in 43. To characterize soil nutrient availability, we collected soil cores at each site and total soil N and P were estimated using the Kjeldahl method and the persulfate digestion method (Carter and Gregorich 2007), respectively, at Soil Analysis Laboratory at Sokoine University of Agriculture, Tanzania. We use a ten-year average for rainfall at these sites that was extracted from the Climate Hazard Group InfraRed Precipitation with Station data (CHIRPS) database (Funk et al. 2015) that provides high resolution precipitation estimates based on both long-term climate averages and weather station data. The different resource gradients are correlated: rainfall-soil N (R=-0.06), rainfall-soil P (R= −0.48), and soil N-soil P (R= 0.62) and therefore, we include all three in the same linear model.

### Estimating herbivory intensity

We defined herbivory intensity (HI) as the ratio of the proportion of plant biomass consumed by herbivores to the total plant biomass produced using calibrated satellite imagery. Briefly, we estimated the total standing biomass at a site from satellite-based vegetation index and predicted the total productivity at a site based on rainfall measures at that location. Rainfall-based productivity was determined by measuring peak seasonal biomass in multiple years inside herbivore fences at seven different sites across the Serengeti, and regressing biomass against rainfall in the previous nine months as estimated from CHIRPS (Funk et al. 2015). Finally, HI for each plot and each year was calculated as 1-(Vegetation index-based biomass/ Rainfall based productivity) averaged across five different sampling years (2001, 2002, 2006, 2009, 2016) (Mohanbabu et al. 2022). The patterns in satellite-based HI along rainfall, N and P gradients are similar to patterns found in grazing intensity in a long-term grazing exclosure experiment (Mohanbabu and Ritchie 2022).

#### Leaf traits

Leaf functional traits were measured in accordance with Pérez-Harguindeguy et al. (2013). At each of the 43 plots, we sampled five individuals of *Solanum* that were at least 5 m apart. For each individual, we collected five fully expanded mature leaves in paper bags and refrigerated them until they could be processed for functional and defense traits. Leaves were then scanned, dried for ~2 weeks at 50°C and weighed. Specific leaf area (SLA) was calculated as the ratio of leaf dry mass to leaf area and is assumed to be a key trait of the leaf economic spectrum (LES). The dried samples were then shipped to Syracuse University for further estimation of leaf nutrient, fiber, and phenolic content.

### Leaf Nutrients

Leaf N and C were estimated using an elemental analyzer (NC2100 CN Analyzer, CE Instruments, Lakewood, NJ, USA) whereas leaf P was estimated using acid digestion followed by malachite green based spectrophotometric estimation. Briefly, 50 mg of ground leaf sample was ashed at 500°C for 5 hours and the residual ash was digested in 5 ml of 6M hydrochloric acid for 30 mins in ceramic crucible. The solution with the dissolved plant residue was filtered with Cellulose Filter Paper (CFP 42) and diluted with distilled water to obtain 100 ml of solution. An aliquot from each sample was then treated with acidified ammonium heptamolybdate and Malachite green in polyvinyl alcohol to obtain a green solution. Absorbance was recorded at 630 nm and the amount of phosphorus was calculated from an absorbance standard curve (Ohno and Zibilske 1991).

#### Defense traits

##### Spine density

For each plant, we recorded the number of spines on a 5cm length of stem at approximately 5cm from the base.

##### Foliar phenolic content

We estimated total foliar phenolic content using the Folin-Ciocalteu assay (Ainsworth and Gillespie 2007). Briefly, 5mg of dried, ground, and sieved leaf sample from each individual was homogenized in 2ml of ice-cold 95% v/v Methanol. After incubating in the dark for 48 hours, the samples were centrifuged at 13,000g for 5 minutes at room temperature. 100ul of the resulting supernatant was aliquoted into a fresh 2ml centrifuge tube, diluted with 100ul of distilled water and 200ul of 10% (v/v) Folin-Ciocalteu reagent (Sigma-Aldrich) was added and mixed thoroughly. After 2 minutes, 800ul of 700mM sodium carbonate solution was added to each tube. The reaction mixture was incubated at room temperature for 20 minutes, before absorbance was recorded at 765nm. The phenolic content is presented as gallic acid equivalents based on a gallic acid standard curve.

##### Leaf structural content

We estimated structural lignin content of *Solanum* leaves using a multi-step sequential digestion with ANKOM 200 Fiber Analyzer (ANKOM Technology, New York USA). The leaf samples for each site were pooled, dried and ground. Approximately 0.5g of the dried leaf material was transferred into special 25u porous bags from ANKOM. The bags were sealed, weighed and treated with neutral detergent solution at 100 °C for 75 min and then washed with hot water to remove any remaining detergents. The bags were washed with acetone to remove excess water and dried at 105 °C overnight. The next day, bags were cooled to room temperature in a vacuum chamber and then weighed, before the process was repeated with acid detergent solution at 100 °C for 60 min. before they were washed with water and acetone, and dried. On the following day, once the bags were cooled and weighed, they were treated with 98% sulfuric acid in a beaker at room temperature for 3 hours with occasional stirring. The bags were then cleaned with hot water and acetone, dried at 105 °C overnight. For the last step, the bag were weighed and ashed in crucibles at 500 °C for 330 min. in a muffle furnace. The difference in the weights between successive steps, specifically after conc. sulfuric acid treatment and ashing were used to estimate the percentage of lignin per dry mass of plant tissue.

### Statistical analyses

For the statistical analyses, we correlated the LES (SLA and Leaf N), and defense traits (spine density, phenolics, and lignin and cellulose). We also included traits that are a measure of plant quality (leaf N:C and leaf P:C) that may affect herbivory levels (Mattson 1980, Schade et al. 2003, Lemoine et al. 2014, Welti et al. 2020, Mohanbabu and Ritchie 2022).

All statistical analyses were performed in R (R Core Team 2020). We estimated Pearson correlation coefficients for each pair of traits using the ‘rcorr’ function package Hmisc (Jr et al. 2021). To explore the relationship between traits of interest in a multivariate space, we ran a principal component analysis (PCA) with ‘prcomp’ in the base package of R. Finally, we also ran a linear regression with lm() to test for the influence of rainfall, soil nutrients and herbivory intensity in driving multivariate trait associations.

## Results

### Bivariate associations between traits

We found that SLA was strongly positively associated with leaf N (Pearson’s R= 0.54, p<0.001) and by extension with leaf N:C (R=0.55; p<0.001), confirming the existence of intraspecific LES in *Solanum* (Fig 2). As expected, all three traits were also negatively associated with lignin and cellulose content (SLA: R=-0.36, p=0.018; Leaf N: R=-0.47, p=0.002; Leaf N:C: R=-0.53, p<0.001). Phenolic content, a chemical defense trait, was significantly negatively correlated with lignin and cellulose (Pearson’s R= −0.34, p=0.024), weakly positively associated with SLA (R=0.26, p=0.098) and uncorrelated with all other traits. Interestingly, spine density which is a key defense trait in *Solanum* was uncorrelated with all other traits including other defense traits. Similarly, leaf P:C also had no significant associations with other traits (Fig 2).

**Figure 2:**
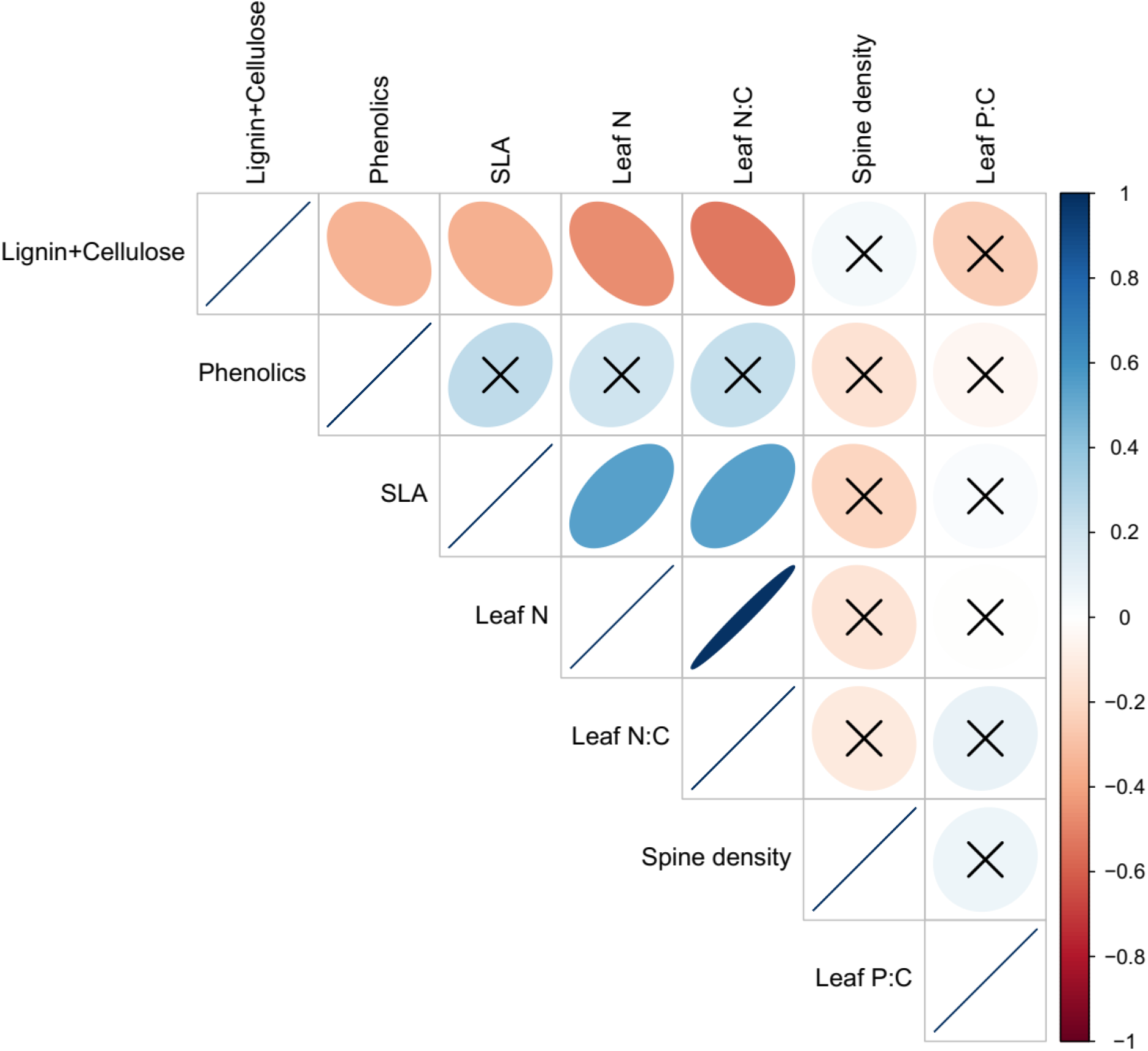
Bivariate associations between traits. The scale from −1 to 1 represents Pearson correlation coefficients and ‘x’ suggests non-significant correlations (i.e., p>0.05).

### Multivariate associations between traits

The first two PC axes explained 59% (72% with 3 axes) of the variation in traits (Fig 3a-c, Table 1). The first component which accounted for 42% of the variation was positively associated with SLA, leaf N, leaf N:C and phenolic content, and negatively associated with lignin and cellulose.

**Figure 3:**
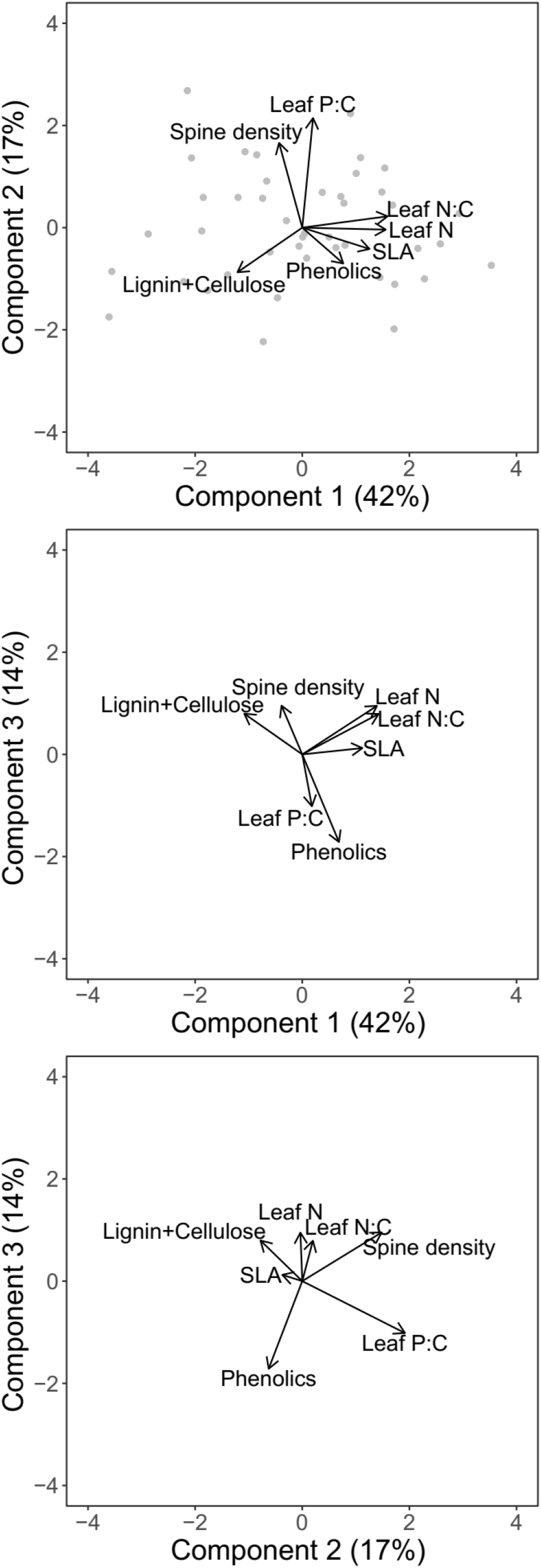
Multivariate associations between traits for different combinations of principal components a) PC1 and PC2; b) PC1 and PC3; and c) PC2 and PC3. The arrows denote the loadings from the PCA and grey point denote sampling sites.

**Table 1:**
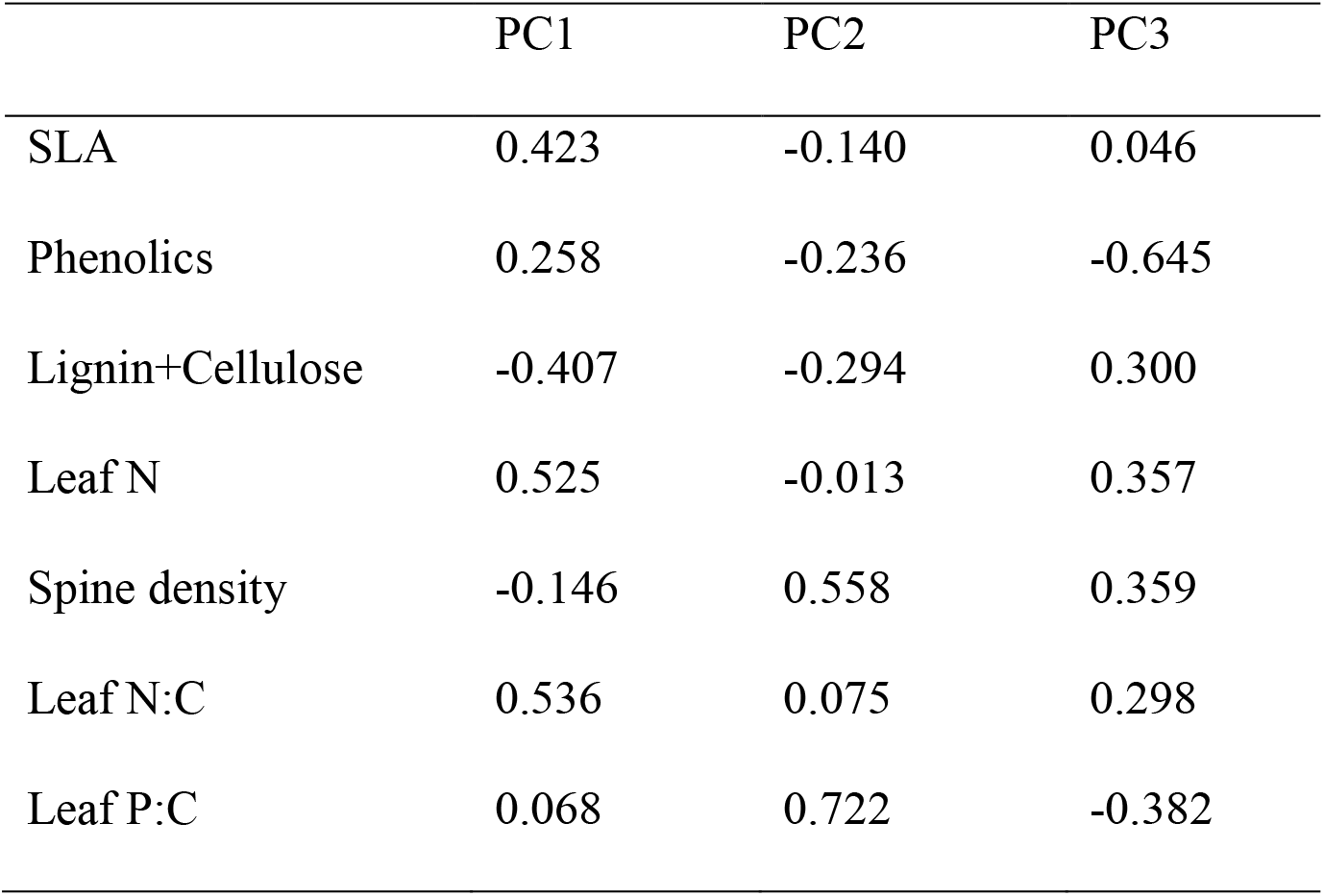
PCA loadings of the different traits for the first three principal components.

The orthogonal component 2 was positively associated with both leaf P:C and spine density and explained 17% of the variation in traits, while component 3 which explained 14% of the variation in traits, was strongly negatively associated with phenolic content. PC axis 3 was also positively associated with leaf N and spine density and negatively with leaf P:C (Fig 3b, c).

### Influence of resource and herbivory intensity on trait associations

Surprisingly, the PC1 axis, which is mostly associated with the leaf economic spectrum was not strongly associated with rainfall, soil nutrients or herbivory intensity (Table 2). Similarly, PC3 was also unassociated with resource and herbivory risk gradients. In contrast, PC2 which is positively associated with physical defenses was positively associated with soil P and herbivory intensity but not rainfall or total soil N.

**Table 2:** Slopes (and standard error) for associations of PC axes with resources and herbivory. Values in bold denote statistical significance at p<0.05 and values in italics denote significant at p<0.1

| Variable | PC1 | PC2 | PC3 |
| --- | --- | --- | --- |
| Resource model |  |  |  |
| Rainfall | -0.49(0.26) | 0.21(0.20) | 0.19(0.19) |
| Total soil N | 0.52(0.36) | -0.16(0.22) | 0.02(0.21) |
| Total soil P | -0.77(0.41) | **0.54(0.25)** | 0.21(0.24) |
| Herbivory model |  |  |  |
| HI | -0.04(0.27) | **0.34(0.16)** | 0.20(0.15) |

## Discussion

In this paper, we provide evidence for intraspecific associations between leaf economic spectrum and defense traits that respond to resource supply and herbivory gradients. As expected, we found positive associations between SLA and leaf N (Fig 1), consistent with predictions of an intraspecific LES (Martin et al. 2017, Fajardo and Siefert 2018). In multivariate trait space, the LES axis described a significant proportion of trait variation and was correlated with lignocellulose at the resource conservative end of the spectrum. Phenolics too showed some positive association with the LES axis but was not strongly correlated with resource acquisitive traits (Fig 2). Spine density was not correlated with the other traits in pairwise comparisons but instead was aligned with leaf P:C in the multivariate space. In contrast to our expectations of resource availability driving the LES axes (Reich et al. 1999, Wright et al. 2004), we found that none of the resources could explain variation in the PC axis 1 which was positively associated with resource acquisitiveness (Table 2). Similarly, variation in PC axis 3 was also not explained by variation in multiple resource supply or herbivory intensity. However, the PC axis 2 which was associated with spine density was also positively associated with soil P and herbivory intensity (Table 2). These results indicate that, 1) resources that influence LES traits may be different from the ones that affect physical defenses; and 2) resource acquisitiveness may independently influence chemical and leaf structural defenses irrespective of resource supply.

Contrary to our initial expectations, we found that the LES axis was unrelated to both the resource and herbivory risk gradients we studied, which may explain the lack of correlation between LES traits and spine density. The intraspecific LES axis was not associated with variation in rainfall or total soil N, two resources that have previously been shown to be important drivers of variation in LES traits (Ordoñez et al. 2009, Dwyer et al. 2014, Maire et al. 2015). Although, LES-resource supply associations are observed even within species, they may be weaker than those for interspecific comparisons (Siefert et al. 2014, Bergholz et al. 2017, Kuppler et al. 2020) and can depend on the study species. Additionally, soil P, a relatively less studied gradient, was also uncorrelated with the LES axis but emerged as the most important resource describing variation in herbivory and spine density across the Serengeti. Such mismatches in the key resource affecting traits are more likely when herbivory risk, i.e., both herbivore abundance and herbivore choice of nutrient rich plant tissue, are included, as they offer additional dimensions of selection pressure on plant traits (Agrawal 2020). Therefore, future attempts to integrate defense traits with other plant functional traits should account for the varying influence of resource supply on plant traits and herbivory risk driving those plant traits.

Regardless of the impact of resources, there was considerable variation in LES traits, and this was associated with allocation of carbon to defenses. For example, our results support the expectation that resource conservative plants with long-lived leaves featuring greater structural C in the form of lignin and cellulose (Fig. 3a) defend against biotic and abiotic agents (Coley 1988, Cabane et al. 2012). This allocation to structural components can pose additional constraints on allocation of C to chemical defenses thereby making it unlikely for resource conservative strategy to also have chemical defenses (Eichenberg et al. 2015). On the other hand, resource acquisitive plants featured greater chemical defenses similar to past studies (Chauvin et al. 2018, Agrawal 2020). Using chemical rather than structural C in defense may be more efficient for resource acquisitive plants because chemical defenses can be constructed or deconstructed over relatively short time scales and thus re-allocated to other functions (Coley et al. 1985). According to the existing hypothesis (Fig. 1), positive associations between the LES axis and herbivory result in correlations between LES and defense traits. However, herbivory intensity and the LES axis (PC1; Fig. 3) were uncorrelated in our study (Table 2), leading us to speculate that LES traits may influence allocation to defenses due to structural constraints, and therefore, independently of herbivory risk.

There was little evidence that LES traits were related to physical defenses, as spine density, the physical defense in our study, was uncorrelated with other defense and LES traits. This result is different from the findings of Armani et al (2020) who found a negative association between spine density and acquisitive growth traits for plants grown in the absence of herbivory. The presence of herbivory in our study may explain some of the mismatch between our results and Armani et al (2020) in which plants did not experience any herbivory during the study. Additionally, it is likely that physical defenses do not directly change leaf function, unlike allocation to lignin or cellulose which can reduce photosynthetic efficiency due to reduced SLA (Poorter et al. 2009). However, spine density was positively associated with mammalian herbivory intensity and soil P hinting that physical defenses may show more consistent associations with risk from herbivory rather than resource acquisitiveness of the plant. Thus, different types of defenses likely dominate the defense-LES trait space, contingent on plant resource strategy and risk from herbivory.

Our results suggest potential for a more complex influence of resource availability and herbivory risk on allocation to different types of defense traits. The current LES-defense framework suggests that structural defense strategy may be favored at low resource acquisitiveness and low herbivory while chemical defense or tolerance strategy may be favored at high resource acquisitiveness (Mason and Donovan 2015, Agrawal 2020, Armani et al. 2020, Morrow et al. 2022). Although herbivory and LES traits were not associated in our study, it is possible that insect herbivory (which we did not measure) is associated with resource acquisitiveness. Hence, we cannot rule out that resource acquisitiveness might produce traits that are favorable to herbivores and consequently increase herbivory risk and associated defenses.

However, expectations for physical defenses remain unclear in the current framework (Fig 4a) (Painter 1951, Agrawal and Fishbein 2006, Mason and Donovan 2015, Agrawal 2020). Therefore, based on our results, we propose a new hypothesis which builds on the existing framework (Fig 1 and Fig 4a) by considering an additional component of herbivory risk that is not directly related to LES traits. This new herbivory component (Herbivory 2 in Fig 4b) may reflect factors that are usually not considered in resource-defense models such as frequency of herbivory (Ritchie and Penner 2020), herbivore dependence on resources not important for plants (e.g., Na) (Borer et al. 2019, Welti et al. 2019, Kaspari 2020) or herbivore vulnerability to predation risk (Anderson et al. 2010, Riginos 2015). In addition, diverse herbivore species assemblages may produce different herbivore response to LES traits: smaller herbivores may prefer resource acquisitive plants with high nutrient content while larger herbivores may consume structurally defended plants as long as they are present in sufficient quantity (Olff et al. 2002).

**Fig 4:**
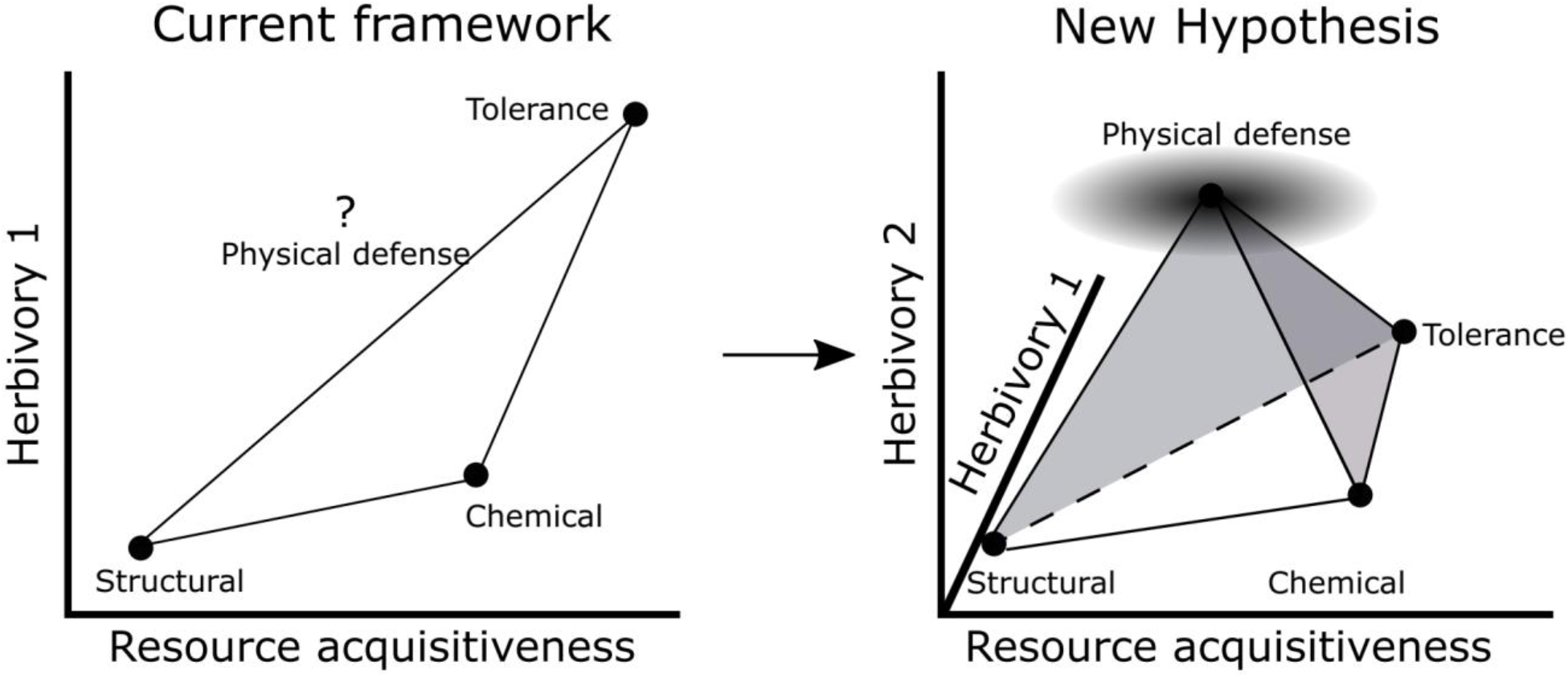
Current and proposed hypotheses to integrate LES and defense traits. a) Current framework which assumes a positive correlation between resource acquisitiveness and herbivory (1) to result in Structural, Chemical and Tolerance strategies. The position of physical defenses is unknown; b) new hypothesis that includes a second axis for herbivory (2) which may depend on frequency of herbivory or other abiotic factors that may not be associated with resource acquisitiveness resulting in Structural, Chemical, Physical defense and Tolerance strategies. The net effect of herbivory is dependent on both Herbivory 1 and 2 axes and may influence the defense strategy of a plant. The shaded gray circle around physical defense suggests that there is potential for large variation in allocation to physical defenses.

The resulting 3D trait space could encompass four different coinciding trait combinations that imply different strategies of response to herbivores: (1) low resource acquisitiveness and low net herbivory risk (structural defense strategy-low SLA, leaf N, and high lignin and cellulose), (2) high resource acquisitiveness and moderate (and infrequent) net herbivory risk likely from insects and/or small mammals (chemical defense strategy-with high SLA, leaf N and phenolics), (3) low (or moderate) resource acquisitiveness and moderate to high herbivory risk (physical defense strategy-low SLA, leaf N, high spine density), and (4) high resource acquisitiveness and high, net herbivory risk (tolerance strategy, i.e., no defense trait) (Fig. 4b). All these scenarios are likely to confer anti-herbivore properties under specific combinations of resource supply and herbivory. Consequently, habitats differing in the range of resource and both components of herbivory may favor different trait combinations in different parts of the tetrahedral trait space. While less parsimonious hypotheses are to be discouraged, we argue that the fourth “point” of the trait tetrahedron is necessary to accommodate the complexities discussed above and provide an explanation for our data. Clearly, further studies, especially experimental ones, are needed to address the independence of herbivory from LES traits across habitats with different resource supply.

While we demonstrate associations between defense and LES traits under natural conditions of resources and herbivory, our study features the inherent limitations of an observational study. The LES traits may be responding to a resource other than rainfall, soil N or P, although based on abundant evidence from other studies on the importance of these specific resources (Wright et al. 2001, 2005, Ordoñez et al. 2009, Moles et al. 2014, Maire et al. 2015), this possibility seems unlikely. Additionally, herbivory on specific *Solanum* individuals may be uncorrelated with the landscape level measures of herbivory intensity. However, plant allocation to defense is often an outcome of herbivory risk experienced over longer time periods of time (Young et al. 2003) and measuring herbivory on each *Solanum* individual over months or years would have been prohibitively expensive. Our measures of herbivory intensity are an average over ten years and thus offer a better representation of long-term herbivory risk experienced at each site.

Furthermore, we are limited in our ability to make causal inferences, but future common garden experiments might shed more light on the processes that drive the patterns between resource-use and defense traits in plants. Finally, our study took advantage of the exceptionally wide range of conditions in which *Solanum* occurs, but patterns in other plant taxa need to be explored to determine how general these results are for intra-specific trait variation.

In summary, we show intraspecific associations between leaf economic spectrum and defense traits along resource supply and herbivory intensity gradients. We found that the LES axis explained a considerable amount of the variation in *Solanum* traits across the Serengeti landscape, but it was unrelated to resource supply or herbivory. In contrast, spine density was positively associated with both soil P and herbivory intensity, suggesting that LES and defense traits could be independently influenced by resources and herbivory. Based on our results, we speculate that different defense traits may become beneficial under different combinations of resource acquisitiveness and herbivory risk and propose a defense tetrahedral model to predict occurrence of different defense strategies. LES strategies have been well-explored for thousands of species globally and integrating defense traits with the LES will provide a multi-dimensional understanding of traits in response to both abiotic and biotic conditions.

## Acknowledgements

We would like to thank Emilian Mayemba for assistance in the field. Special thanks to TAWIRI, COSTECH, and TANAPA for allowing us to carry out research at Serengeti National Park. The study was supported by NSF grant DEB 1557085. The authors have no conflict of interest to declare.

## Author contributions

NM and MR planned, designed, and interpreted the research. NM collected all the field and trait data with help from DJ for lab-based analyses while MV contributed to estimating herbivory intensity for the sites. NM wrote the first draft of the manuscript and NM, MR and MV contributed substantially to revising the manuscript.

## Data Availability

All data will be made available on Dryad upon acceptance of the manuscript

